# A scalable computational framework for annotation-independent combinatorial target identification in scRNA-seq databases

**DOI:** 10.1101/2024.06.24.600275

**Authors:** Maximilian Buser, Carsten Marr

## Abstract

The applicability of novel targeted cell therapies is constrained by on-target off-tumor toxicities, posing significant challenges for safe and effective treatment. To develop strategies for mitigating these toxicities, a comprehensive understanding of the distribution of single-cell profiles across human tissues is essential. Single-cell RNA sequencing (scRNA-seq) provides detailed insights into cellular diversity and function. However, as data collection expands and RNA expression atlases comprise tens of millions of cells, effectively querying these databases to identify cell populations matching specific gene expression patterns becomes challenging. Key challenges include managing data volume, handling batch effects, navigating study-dependent data curation and annotation, and addressing the combinatorial complexity of possible target patterns. Userfriendly screening tools that allow for complex query target patterns without relying on existing annotations are currently missing. Here we present `otopia`, a computational framework that enables fast queries of large scRNA-seq databases, enhancing the precision of cell population identification. Using precomputed neighborhood graphs and boolean logic for expression patterns, `otopia` enables efficient queries addressing arbitrary combinatorial target patterns. `otopia` does not depend on cell type annotations and by exploiting unbiased expression pattern matching scores, it uncovers critical populations that annotation based methods overlook. We exemplify `otopia`’s capabilities by identifying the largest B cell populations in the CELLxGENE census data using a CD19^+^CD22^+^CD5-query target pattern. Moreover, we show that `otopia` can provide a concise overview of cell populations matching hypothetical acute myeloid leukemia (AML) dual targets, realizing logic AND gates, across different tissues. `otopia` enables the estimation of ontarget off-tumor toxicities for drugs targeting both single markers and complex marker combinations, surpassing previous methods that focus mainly on single markers. With the expanding availability of scRNA-seq data, `otopia` represents a scalable, unbiased, and automated approach to refine putative therapeutic targets in order to minimize toxicities.

## Introduction

Comprehensive single-cell multi-omics datasets, combined with computational analysis methods, have become essential for the rational design of chimeric antigen receptor T cell [1–3] and other antibody-based targeted therapies [4–6]: The computational screening of malignant, tumor-specific reference data [7, 8] facilitates the identification of efficacious targets [9–13]. Complementarily, the screening of healthy reference data [14–16] supports safety evaluation by predicting on-target off-tumor toxicities in non-malignant cells [17], thereby guiding the development of strategies to mitigate such toxicities [18, 19]. However, current approaches for the prediction of toxicities from transcriptomic data face several challenges: (i) Study-dependent curation and cell-type annotations may not necessarily be comparable between datasets and may not fit the resolution required to faithfully estimate toxicities. (ii) Custom reference datasets employed for the computational estimation of toxicities and efficacies of drug candidates are difficult to maintain, leading to data silos that are not further utilized. (iii) Combinatorial target patterns promise sensitivity and specificity boosts, but are hard to precompute in advance due to their abounding number. (iv) There is a lack of suitable metrics that can efficiently cope with batch effects across different datasets and combinatorial target patterns. Computational approaches that help to overcome these problems and can readily utilize public, community-driven, and version-tagged databases are thus needed to advance drug design. Here we present otopia, a computational framework that enables efficient queries of combinatorial target patterns in large scRNA-seq databases, enhancing the identification of critical cell populations that are *on target*.

## Results

### Identifying populations matching combinatorial targets in single-cell data

To identify cell populations matching gene expression patterns in an unbiased manner and independent of existing single-cell annotations, otopia uses a precomputed neighborhood graph (Fig. 1A). Each vertex of this graph represents a single cell, and the graph collectively accounts for all the cells in the reference data. Depending on the specific question, a suitable partitioning of the reference data might be necessary. The neighborhood graph can be efficiently precomputed from dimensionally reduced representations of the single-cell reference profiles.

**Figure 1:**
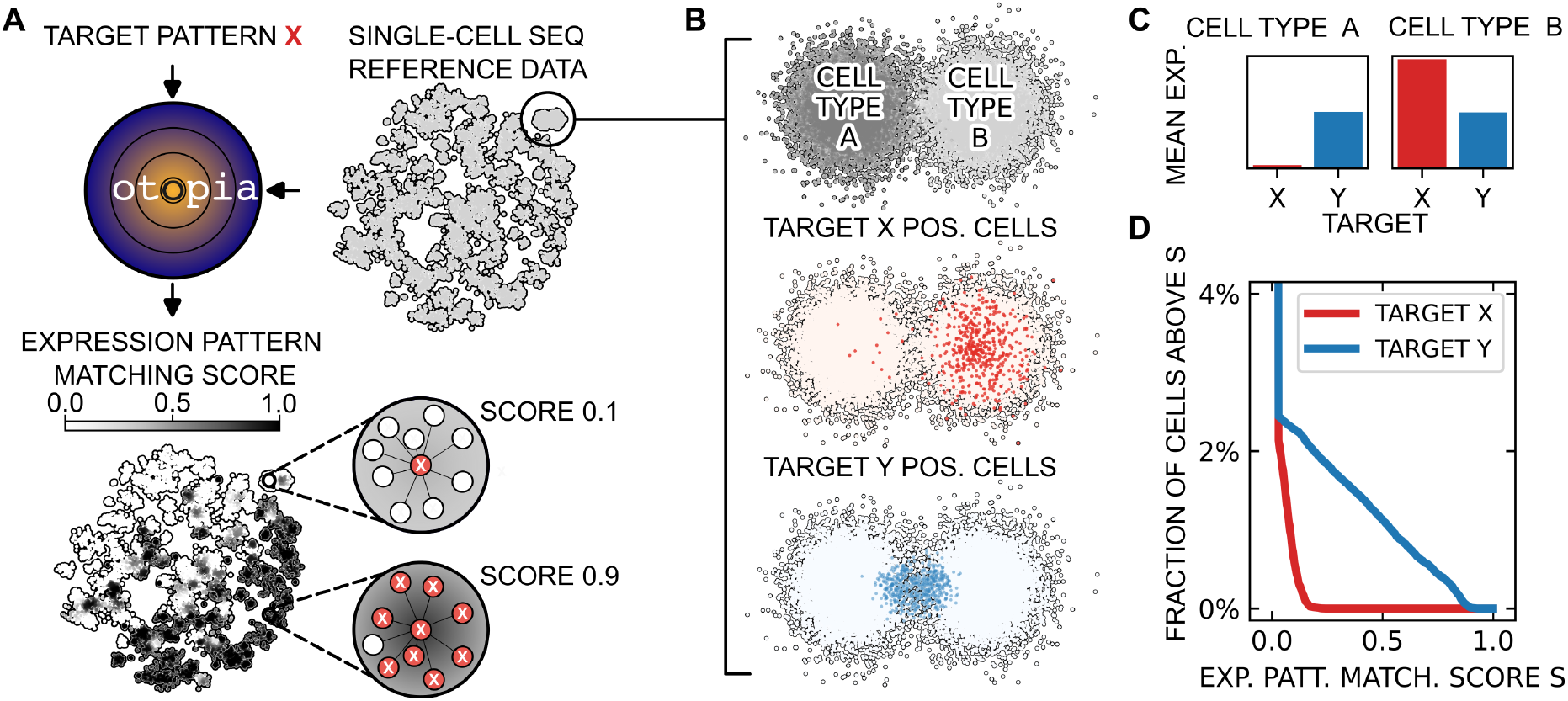
otopia identifies cell populations matching query target patterns in large single-cell reference data. (**A**) otopia screens large single-cell sequencing reference databases, such as the CELLxGENE census, to identify populations that match query target patterns without relying on existing annotations. To enable efficient queries, otopia uses precomputed neighborhood graphs to calculate expression pattern matching scores at the single-cell level. The expression pattern matching score of a cell is defined as the fraction of cells among its *K*-nearest neighbors that match the pattern, exemplified here for *K =* 9. Irrespective of a cell’s neighborhood, its score is set to zero if it does not match the target profile itself. (**B**) Synthetic data illustrating the neighborhood-based approach underlying otopia. Single cells are shown in a two-dimensional embedding revealing two major clusters with fictitious cell-type annotations A and B and the expression of target pattern X and Y. Target pattern X-positive cells are sparsely but homogeneously expressed across cell type B, whereas Y-positive cells are found in a dense subcluster at the borderline between A and B. (**C**) A standard target pattern analysis on the average expression per cell-type level misses the dense subcluster of target pattern Y-positive cells. (**D**) The expression pattern matching score distributions for X and Y, which account for all cells from A and B, indicate the presence of a sparse target pattern X-positive population and a dense target pattern Y-positive population.

With otopia, we can consider queries for arbitrary combinatorial expression patterns. For each single or combinatorial target, we calculate an expression pattern matching score for every cell in the reference data (Fig. 1A). Specifically, the score of a cell is defined as the fraction of cells among its *K*-nearest neighbors that match the target pattern. If a cell does not match the target pattern, its score is set to zero, regardless of its neighborhood. By accounting for the individual neighborhood of single cells, the score continuously reflects the density of cells matching the target pattern across subpopulations.

Compared to annotation based methods, otopia’s approach efficiently uncovers critical populations that do not necessarily correlate with coarse-grained clusters and cell-type labels (Fig. 1B-D).

### CD19^+^CD22^+^CD5^-^ query reveals largest B cell populations among 23 million cells

We exemplify otopia’s capability by querying 23,793,940 human cells, which have been provided to the CELLxGENE census [16], for populations matching the B cell expression pattern CD19^+^CD22^+^CD5^-^ (Fig. 2A). Specifically, we have filtered the CELLxGENE census (v.2023-12-15) for non-pathological cells and 4, 443 unique tissue-dataset-donor combinations with at least 100 cells each have been considered for downstream analyses. Raw single-cell profiles corresponding to each of these combinations have been normalized to 10, 000 counts per cell and log-transformed after the addition of one. A dimensionally reduced representation of these profiles has been obtained by accounting for the top 50 principal components of the top 2, 000 highly variable genes per combination [23, 24]. We have used PyNNDescent [25] to compute nearest neighbor graphs for each tissue-dataset-donor combination based on these reduced representations.

**Figure 2:**
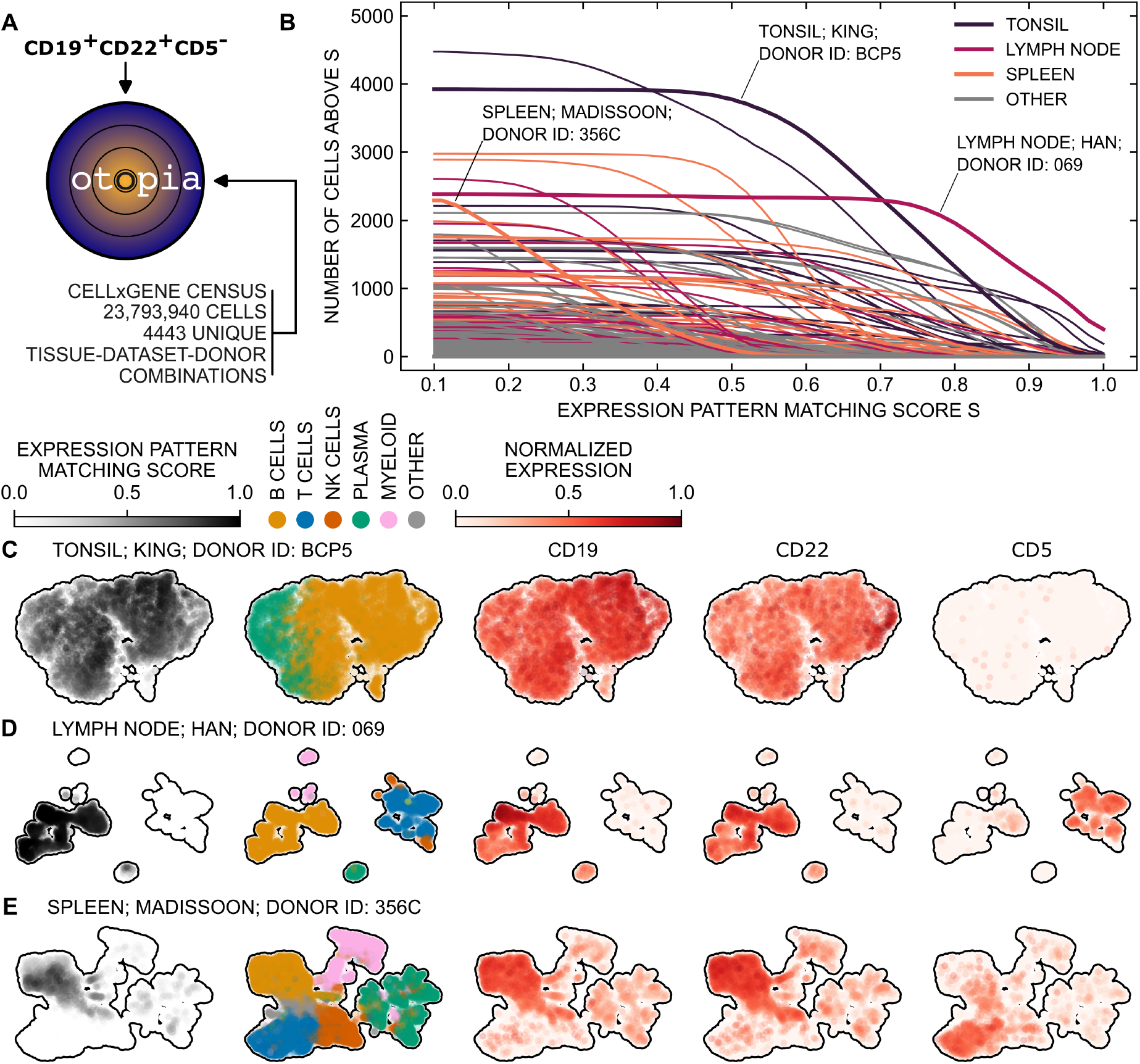
otopia identifies the largest B cell populations in CELLxGENE census data based on a CD19^+^CD22^+^CD5^-^ query target pattern. (**A**) We query the CELLxGENE census for cells with a neighborhood matching the expression pattern CD19^+^CD22^+^CD5^-^, which is typical for B cells. (**B**) optopia identifies tissues with large B cell fractions. The lymph node sample from donor 069 from the dataset provided by Han et al. [20] comprises the largest number of cells with an expression pattern matching score above 0.75. Also the tonsil sample from donor BCP5 provided by King et al. [21] and the spleen sample from donor 356C provided by Madissoon et al. [22] show high scores. (**C-E**) Transcriptomic profiles of all cells from these samples are projected into two-dimensional UMAP embeddings. The UMAPs show the gene expression pattern matching scores, coarse-grained cell type annotations as detailed on in the main text, and the normalized expression levels for CD19, CD22, and CD5. The fractions of CD19^+^CD22^+^CD5^-^ B cells in the tonsil, lymph, and spleen samples are 56%, 84%, and 14%, respectively.

An analysis of the number of cells exceeding various thresholds of the CD19^+^CD22^+^CD5^-^ expression pattern matching score shows that tonsil, lymph node, and spleen samples in the reference data contain large cell populations matching the query pattern (Fig. 2B). Note that here and throughout the following examples, we consider *K =* 50 nearest neighbors for the calculation of expression pattern matching scores. Resolving the data from three exemplary datasets [20–22] for expression pattern matching scores, cell-type annotations, and normalized expression levels in two-dimensional UMAP embeddings reveals that cells with high scores are labeled as B cells in the CELLxGENE census (Fig. 2C-E). For clarity, we use coarse-grained cell-type annotations in Fig. 2C-E: B cells include follicular B cells and memory B cells; myeloid cells include conventional dendritic cells, dendritic cells, macrophages, monocytes, and plasmacytoid dendritic cells; and plasma cells include IgG plasma cells, IgM plasma cells, and plasmablasts.

### Inter-tissue comparison of cell populations matching putative AML dual targets

Using the same reference data comprising 23, 793, 940 cells, we turn to address six putative AML single targets: CD7^+^, HAVCR2^+^, IL3RA^+^, CD33^+^, CLEC12A^+^, and CD244^+^, as well as all fifteen dual combinations (CD7^+^HAVCR2^+^, CD7^+^IL3RA^+^, …), implementing logic AND gates for pairs of these targets [10, 26, 27]. An overview of the fraction and number of cells with a gene expression pattern matching score of at least 0.2 for different tissues reveals that, in addition to blood and bone marrow, which are expected to rank highly, lung and spleen samples show high scores for many of the dual targets (Fig. 3A). For clarity, the tissues considered here correspond to the general tissue labels of the CELLxGENE census [16], and tissues with fewer than 1, 000 cells exceeding a score of 0.2 for any of the dual targets have been omitted.

**Figure 3:**
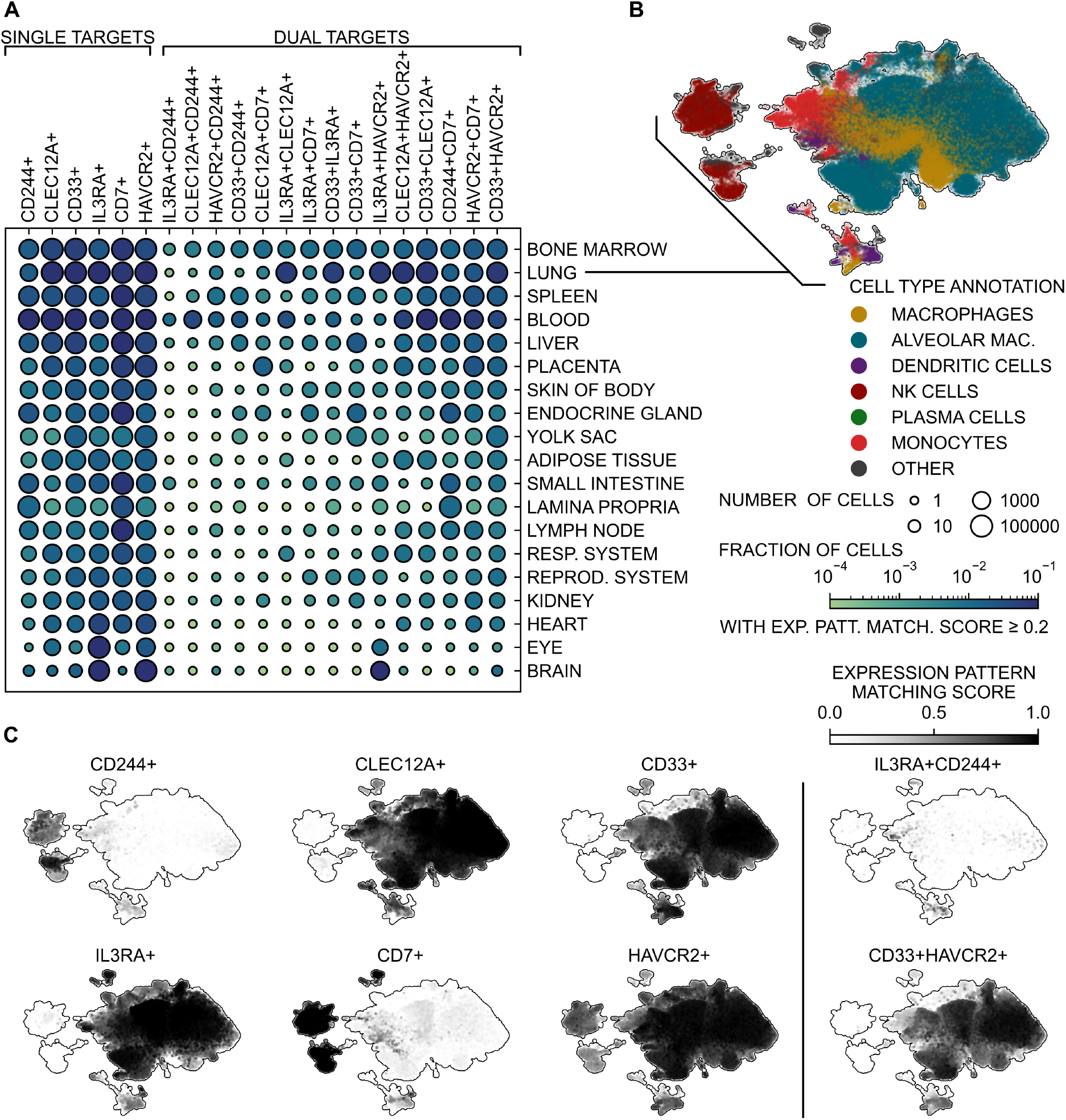
Addressing six hypothetical AML single targets (CD7^+^, HAVCR2^+^, IL3RA^+^, CD33^+^, CLEC12A^+^, and CD244^+^) and 15 dual combinations with otopia reveals that the highest fraction of cells matching most of the dual target patterns are found in the lung and spleen, preceded by bone marrow and followed by blood. (**A**) Fraction and number of cells with a gene expression pattern matching score of at least 0.2 in different tissues. For visualization purposes, the list of tissues considered here is restricted to those with at least 1, 000 cells reaching a score above 0.2 for at least one of the dual targets. (**B**) 143, 220 lung cells with a gene expression pattern matching score of at least 0.2 in at least one of the 15 dual targets are resolved in a joint two-dimensional UMAP embedding with their annotated cell type. (**C**) In the same embedding, the expression pattern matching scores assigned to each of these lung cells are shown for all single targets as well as for CD33^+^HAVCR2^+^ and IL3RA^+^CD244^+^.

There are 143, 220 lung cells with an expression pattern matching score above 0.2 for at least one of the dual targets. This critical population spans cells from 135 different donors and 15 different studies, with 72% of the cells corresponding to the datasets provided in Refs. [28–30]. A two-dimensional UMAP representation based on a 50-dimensional scvi embedding [31] of the underlying single-cell profiles resolves the originally annotated cell types (Fig. 3B).

A visualization of the expression pattern matching scores for the dual targets IL3RA^+^CD244^+^ and CD33^+^HAVCR2^+^ in the same UMAP embedding highlights only a limited number of cells matching IL3RA^+^CD244^+^, whereas CD33^+^HAVCR2^+^ is prevalent, particularly among macrophages (Fig. 3C). This observation aligns with scores for the single targets: NK cells and macrophages exhibit contrasting expression patterns with NK cells showing high scores for CD244^+^ but not for IL3RA^+^, and macrophages displaying high scores for IL3RA^+^ and HAVCR2^+^.

## Discussion

We introduced otopia, a computational framework designed for efficiently querying large-scale scRNA-seq databases to identify cell populations matching single targets, as well as complex combinatorial gene expression patterns. This capability is crucial for understanding and mitigating on-target off-tumor toxicities in targeted cell therapies. Unlike annotation-based approaches, otopia uses precomputed neighborhood graphs and a novel gene expression matching score to identify critical cell populations in an unbiased manner.

Our analysis with otopia, utilizing 23, 793, 940 reference cells, demonstrates its effectiveness in handling large-scale data. Specifically, otopia successfully identified B cell populations matching a CD19^+^CD22^+^CD5^-^ query pattern and estimated toxicities associated with putative AML dual-targets. Moreover, our newly introduced score paves the way for concise overviews and automated analyses: By considering binned distributions of the score and by marginalizing meta-data categories, such as dataset, tissue, donor, or disease labels, results concerning large-scale reference databases can be efficiently condensed. Thus, otopia successfully overcomes the challenges outlined at the beginning, including managing data volume, handling batch effects, navigating study-dependent data curation and annotation, and addressing the combinatorial complexity of target patterns.

Turning to the limitations of the computational framework presented here, it is important to emphasize that while otopia effectively addresses toxicities, it does not evaluate the efficacy of putative targets through malignant reference data screening. This underscores the necessity of integrating otopia with complementary methods that predict the efficacy of therapeutic targets [9–11, 13]. Furthermore, although the robustness of prediction results can be evaluated methodically, for instance, by tuning the hyperparameter *K*, faithful predictions of on-target off-tumor toxicities demand rigorous and experimental validation to ensure clinical relevance and safety [12, 18].

We anticipate that otopia’s data-driven, unbiased, and scalable approach will aid in refining therapeutic targets for antibody-based cell therapies, advancing precision medicine by utilizing large-scale scRNA-seq reference datasets. The code will be made available at github.com/marrlab.

## Acknowledgments

We thank Stephanie Frenz-Wießner for valuable comments and discussions. CM has received funding from the European Research Council (ERC) under the European Union’s Horizon 2020 research and innovation program (Grant agreement No. 866411).

## Notes

### Competing Interest Statement

The authors have declared no competing interest.

